# Taguchi–Machine Learning Hybrid Framework for Optimization of Particulate Drug Delivery Systems

**DOI:** 10.1101/2025.08.16.670654

**Authors:** Doğukan Duymaz, Seda Kizilel

## Abstract

Optimizing particulate drug carrier systems requires balancing multiple formulation parameters to achieve target physicochemical properties while minimizing experimental burden. Here, we implement a hybrid optimization framework integrating a Taguchi orthogonal array (OA) design with statistical modeling and machine learning (ML)–based interpretability, using doxorubicin-loaded chitosan microspheres (DOX-CS MSs). Microspheres were synthesized via a water-in-oil emulsion crosslinking method and characterized by FTIR, XRD, and FESEM to confirm chemical structure, crystallinity, and spherical morphology. The optimization targeted a particle size of 5–7 µm and encapsulation efficiency (EE) >90%. An initial L₉ Taguchi OA design efficiently narrowed the formulation space by varying chitosan concentration (1–3% w/v), glutaraldehyde concentration (1.5–5% v/v), and crosslinking time (3–5 h), yielding nine core formulations. Pearson/Spearman correlation, second-order polynomial regression (Poly²), and Gradient Boosting Machine (GBM) models quantified parameter influences and predicted performance. SHapley Additive exPlanations (SHAP) identified chitosan concentration as the primary determinant of both size and EE, with glutaraldehyde content exerting secondary, synergistic effects. Poly² response-surface modeling achieved high predictive accuracy (R² = 0.983 for size; R² = 0.986 for EE) and yielded explicit regression equations for real-time formulation targeting. This hybrid Taguchi–ML approach enables rapid factor prioritization, reveals nonlinear interactions overlooked by conventional Taguchi analysis, and offers transparent ML interpretability. Beyond chitosan-based carriers, it provides a generalizable, scalable route to rational formulation design in complex particulate systems for targeted biomedical applications.

## 1. INTRODUCTION

Micro- and nanoparticle systems have emerged as versatile platforms with extensive applications across various biomedical domains, including controlled drug delivery, targeted therapeutics, diagnostics, tissue engineering, regenerative medicine, and wound healing. The ability of these particulate systems to precisely regulate the spatial and temporal release of bioactive agents, control cellular interactions, and modulate biological responses positions them as invaluable tools in modern biomedical research ^1–4^. Despite their promising potential, successful translation from bench to bedside remains challenging due to stringent requirements for reproducibility, precise size distribution, efficient encapsulation, and predictable performance in biological environments.

Polymeric particulate systems, particularly those fabricated from biodegradable polymers, have gained significant attention due to their intrinsic biocompatibility, tunable physicochemical properties, and predictable biodegradation profiles. Among biodegradable polymers, chitosan (CS) has been widely explored owing to its natural origin, inherent biocompatibility, biodegradability, low immunogenicity, and intrinsic antimicrobial and immunomodulatory properties, making it suitable for various biomedical applications ranging from drug delivery systems to tissue scaffolds ^5, 6^. Importantly, chitosan-based particulate systems offer extensive modifiability, enabling tailored physicochemical properties such as particle size, surface charge, and drug encapsulation profiles. For instance, CS microparticles encapsulating doxorubicin (DOX), a widely used model chemotherapeutic drug, have demonstrated significant promise due to their enhanced stability, sustained drug release characteristics, and improved therapeutic outcomes in targeted cancer therapies ^7^. DOX, as a model drug, is particularly useful because of its extensive characterization in various drug delivery systems, well-documented biological activities, and relevance to numerous clinical scenarios ^8^.

In the context of particulate drug delivery systems, particle size is a critical determinant of cellular uptake, biodistribution, and overall therapeutic efficacy ^9, 10^. Tailoring the size of particles can enable selective interactions with specific cell populations, phagocytic cells such as macrophages preferentially internalize particles in the 1–3 µm range, whereas smaller nanoparticles may escape immune recognition or accumulate passively at disease sites via enhanced permeability and retention (EPR) effects ^11–13^. Thus, defining an optimal size window based on application-specific biological interactions is essential for engineering efficient particulate carriers. Alongside size, encapsulation efficiency (EE) remains a vital parameter in micro/nanoparticle design, as high EE ensures effective drug loading, minimizes therapeutic agent loss, and improves dosing precision, especially in applications involving costly or potent compounds.

Optimizing these critical properties typically requires extensive experimentation and meticulous control over synthesis conditions. Conventionally, optimization studies rely on the “one-factor-at-a-time” (OFAT) approach, which not only increases experimental burden but also fails to accurately capture interactions between multiple factors ^14^. To address this challenge, statistical design of experiment (DoE) type methods, such as the Taguchi method, are increasingly employed due to their efficiency in systematically exploring multiple experimental parameters with significantly reduced experimental burden ^15–18^. The Taguchi method, pioneered by Genichi Taguchi, is a robust statistical approach designed to optimize process parameters and enhance product quality while minimizing experimental effort. It leverages orthogonal array (OA) designs to systematically evaluate the influence of multiple factors and their levels on a target outcome. Unlike traditional full-factorial designs that grow exponentially with more factors, Taguchi’s OA-based layout requires only a fraction of the total experimental combinations, drastically reducing cost, time, and labor. Each OA ensures that the influence of each factor is independently and efficiently assessed, while maintaining statistical balance and orthogonality ^19, 20^.

A key advantage of the Taguchi method is the incorporation of Signal-to-Noise (S/N) ratios, which quantify both the mean performance and variability of the system, promoting robust optimization under real-world uncertainty. The method categorizes quality objectives (e.g., “larger-the-better,” “smaller-the-better,” or “nominal-the-best”) and applies corresponding S/N ratio formulas to enhance both performance and consistency ^19, 21^. In biomedical applications, including the optimization of particulate synthesis, the Taguchi method has been effectively used to evaluate the impact of variables such as polymer concentration, crosslinker dosage, reaction time, molar ratio of solvents, temperature and etc. Recent studies have specifically applied this method to optimize drug encapsulation, nanoparticle fabrication, and scaffold design, demonstrating its utility in reducing experimental burden without compromising scientific rigor ^22–27^. However, despite its experimental efficiency, the Taguchi method suffers from several interpretational limitations, particularly when applied to complex biomedical material systems: First of all, Taguchi designs typically assume linearity between factors and outcomes, potentially leading to inaccurate modeling in nonlinear systems ^20^; secondly they offer limited capabilities in detecting main effects and interactions, thus hindering mechanistic insights ^28^; finally, they lack statistical rigor in terms of hypothesis testing and model validation, which can compromise reproducibility and confidence in findings ^17^. In addition, for systems with numerous interdependent variables, Taguchi’s OA structure may oversimplify the experimental space, neglecting higher-order effects ^28^.

To overcome these interpretational limitations, data-driven approaches and computational modeling techniques such as machine learning (ML) algorithms can be integrated with statistical designs. ML methodologies facilitate a comprehensive understanding of complex datasets, allowing precise modeling and prediction of particulate characteristics based on experimental inputs. Such approaches significantly enhance the ability to interpret experimental data, reveal hidden interactions among factors, and provide robust predictive models that enable precise control over particle or, in general, biomaterial properties ^29–31^.

Thus, in this study, we hypothesize that combining the Taguchi experimental design method with ML-driven analyses can systematically optimize microparticle properties, enabling highly predictable and reproducible outcomes. We employ chitosan microspheres (CS MSs) encapsulating the model drug DOX, as a representative biomedical system, to target a desired particle size range (5–7 µm) with high EE (>90%). Our four-step integrated workflow comprises; **(i)** selection of factors and levels for a Taguchi design; **(ii)** synthesizing and characterizing the orthogonal-array formulations; **(iii)** performing comprehensive ML-driven statistical analyses; **(iv)** and generating response-surface models for prediction (**Figure 1**). CS concentration, glutaraldehyde (GA) crosslinker level, and crosslinking time were selected as factors and each was varied at three levels in an L₉ Taguchi array to minimize the number of experiments. DOX-encapsulating microspheres were synthesized via a water-in-oil (w/o) emulsion and characterized for particle size, EE and drug loading, after which multiple computational analyses were performed for rigorous data interpretation.

**Figure 1.**
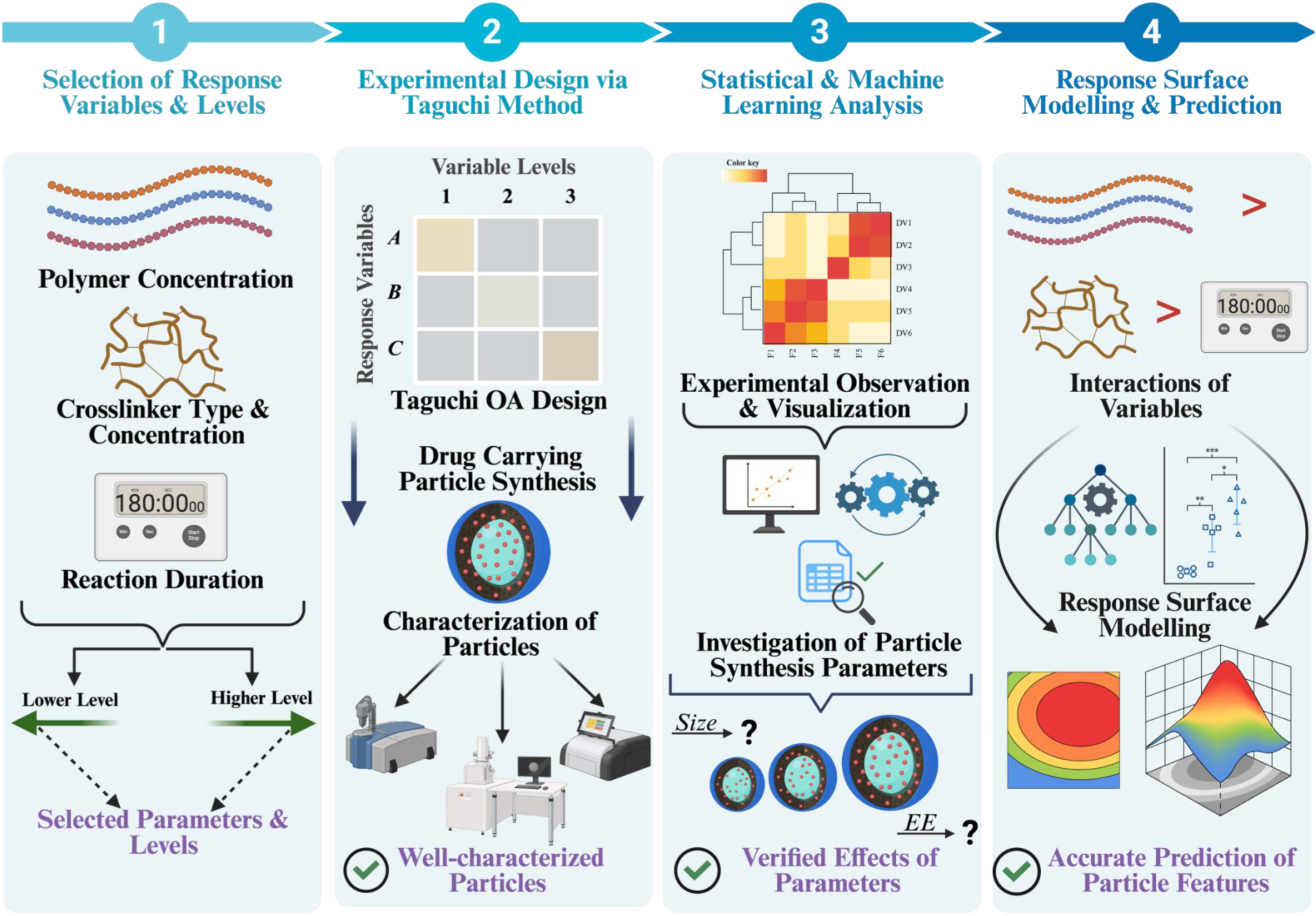
Overview of our four-step design-to-prediction workflow: (**1**) selection of synthesis factors and levels, (**2**) Taguchi experimental design and particle characterization, (**3**) statistical and machine-learning analysis of factor effects, and (**4**) response-surface modeling for formulation optimization. Created in https://BioRender.com.

Our Taguchi S/N-ratio screening and Pearson/Spearman correlation analyses first pinpointed CS concentration as the dominant factor governing both particle size and EE. Subsequent ML modeling, using both Gradient Boosting Machine (GBM) and second-order polynomial regression (Poly^2^), was rigorously validated by leave-one-out cross-validation (LOOCV), yielding promising LOOCV-R^2^ values of 0.89 for size and 0.90 for EE with Poly^2^ model. Permutation importance and SHAP analyses quantitatively confirmed CS% as the top driver, with GA concentration and reaction time playing secondary, synergistic roles. The resulting quadratic model delivers explicit predictive equations (R^2^ = 0.983 for size; R^2^ = 0.986 for EE) that allow rapid contour and 3D surface maps for formulation optimization. Altogether, this concise Taguchi–ML hybrid framework not only minimizes experimental burden but also produces highly accurate, interpretable response surfaces, laying a versatile foundation for tailoring particulate biomaterials across diverse biomedical applications.

## 2. EXPERIMENTAL SECTION

### 2.1 Chemicals and Reagents

All materials used in this study have been obtained from Sigma Aldrich unless otherwise stated.

### 2.2 Preparation of Chitosan Microspheres

Chitosan microspheres were fabricated using an improved emulsion cross-linking method exemplified by several papers in literature ^23, 32^. Initially, a chitosan solution was prepared by dissolving chitosan (Sigma Aldrich, 448869) in a 2% (v/v) aqueous glacial acetic acid (Sigma Aldrich, A6283) solution through magnetic stirring overnight. Subsequently, 3 mL of this chitosan solution was injected into 20 mL of mineral oil (Sigma Aldrich, 330779) containing Span 80 (1%, v/v) (Sigma Aldrich, S6760) using a 21-gauge syringe. This resulted in the formation of a water-in-oil (w/o) emulsion, achieved by vigorously stirring the two phases at 1000 RPM using a mechanical stirrer at room temperature. After 30 minutes of high-shear homogenization, GA (Sigma Aldrich, 354400) was gradually added via a 27-gauge syringe to initiate cross-linking and stabilize the particles. The resulting microspheres were isolated by centrifugation at 4000 RPM and washed thrice with hexane (Sigma Aldrich, 139386) and acetone (Sigma Aldrich, 179973), respectively. The final microsphere sediment was air-dried at room temperature and stored at 2-8°C for subsequent use.

To incorporate a drug into the CS MSs, a desired amount of the drug (∼2.5 mg/mL in total w/o solution) was integrated into the 3 mL CS solution, followed by thorough mixing. This drug-loaded chitosan solution was then emulsified in the same mineral oil phase to form a w/o emulsion. The subsequent steps, including cross-linking, separation, washing, and drying, were identical to the protocol for producing drug-free CS MSs. DOX (Sigma Aldrich, PHR1789) was preferred as a model drug for CS MS studies. All formulations were prepared in at least two independent replicates.

### 2.3 Encapsulation Efficiency of Drug-Loaded Chitosan Microspheres

As the DOX was the model drug utilized for the CS MSs, the absorbance value of 0.01, 0.02, 0.04, 0.08, 0.12, 0.16 and 0.2 mg/ml of DOX were measured at 480 nm using a plate reader to generate a standard curve in the beginning of the experiment (**Figure S1**). EE quantifies the proportion of the actual drug encapsulated within the microparticles relative to the initial theoretical drug amount added during synthesis. Drug Loading (DL) represents the percentage of the microparticle’s weight composed of the loaded drug. To be able to accurately measure both EE and DL of formulations of DOX-loaded chitosan microspheres, the supernatant of centrifuged microspheres in each step of washing was collected and stored for analyzing the free and non-encapsulated DOX during the microsphere formation. EE and DL were calculated with the equations demonstrated below ^33^ (n=2):

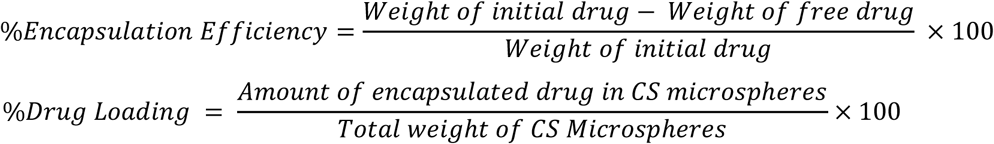

### 2.4 Characterization of Chitosan Microspheres

*FTIR Analysis.* Fourier-transform infrared spectroscopy (FTIR, Nicolet iS10 instrument, ThermoFisher, United States) was employed to analyze the chemical composition of doxorubicin, chitosan and chitosan microspheres with a resolution of 4 cm^−1^. By examining the absorption peaks and frequencies of average of 64 scans in the infrared spectra, covering a range of 650 to 4000 cm^−1^, the specific molecular groups present within these biomaterials were identified. In short, FTIR spectroscopy was utilized to verify the successful incorporation of doxorubicin, into the CS matrix, and formation of microspheres by aiming to identify any chemical differences between the produced microspheres.

*Morphological Analysis.* Size, porosity, and surface morphology of the produced chitosan microspheres were visualized using a ZEISS Ultra Plus Field Emission Scanning Electron Microscope (FE-SEM, ZEISS, Oberkochen, Germany). Before SEM analysis, CS MSs were air-dried completely. Additionally, all the samples were gold-sprayed before FE-SEM scanning.

*XRD Analysis.* X-ray diffraction (XRD) analysis was employed to investigate the crystalline or amorphous nature of microsphere samples before and after synthesis. Additionally, XRD was utilized to confirm the formation of microspheres and encapsulation of doxorubicin within the chitosan matrix. XRD patterns were acquired using a Bruker D2 Phaser X-Ray Diffractometer equipped with graphite-filtered Cu Kα radiation (λ = 1.5418 Å) operating at 40 kV and 20 mA. Data was collected over a 2θ range of 5° to 80° with a step size of 0.021° and a scan rate of 0.8 s per step.

### 2.5 Determination of Mean Size of Microspheres

The synthesized empty and DOX-loaded chitosan microspheres’ sizes (≥100 particles) were analyzed by investigating SEM images of produced particles through Image J software. Obtained size data for each condition were further processed with frequency and gaussian distribution models to approximately calculate the mean size of the produced microspheres for both Taguchi orthogonal array setup and other incorporated conditions. All formulations were prepared in at least two independent replicates to assess reproducibility and quantify experimental variability.

### 2.6 Experimental Optimization of Microspheres by Taguchi Orthogonal Method

Taguchi orthogonal method was applied to evaluate the influence of CS concentration (X_1_), cross-linking time (X_2_), and crosslinker (GA) concentration (X_3_) on the microspheres’ properties (**Table 1**). The designed system was composed of 3 variables set at 3-levels of each parameter, where the mean particle size and drug EE were the main investigated dependent variables. In this system, according to preliminary results of the optimization of desired CS MSs, we have selected 3 main conditions with each CS concentration, which had the highest possibility to give the desired range of size of microspheres, high EE and acceptable DL percentage according to our hypothesis. Other parameters have been defined with respect to Taguchi orthogonal method, constructing an L_9_ orthogonal matrix with most promising conditions, to see the effect of these 3 different parameters varying with each other. While the L_9_ array efficiently screens the main effects of CS concentration, GA concentration, and crosslinking time, it cannot fully capture curvature and two-way interactions, therefore, several of further experimental formulations (shown as black-colored runs in **Table 1**) were also included to fully capture curvature and two-way interactions of these effects accurately and consistently.

**Table 1.**
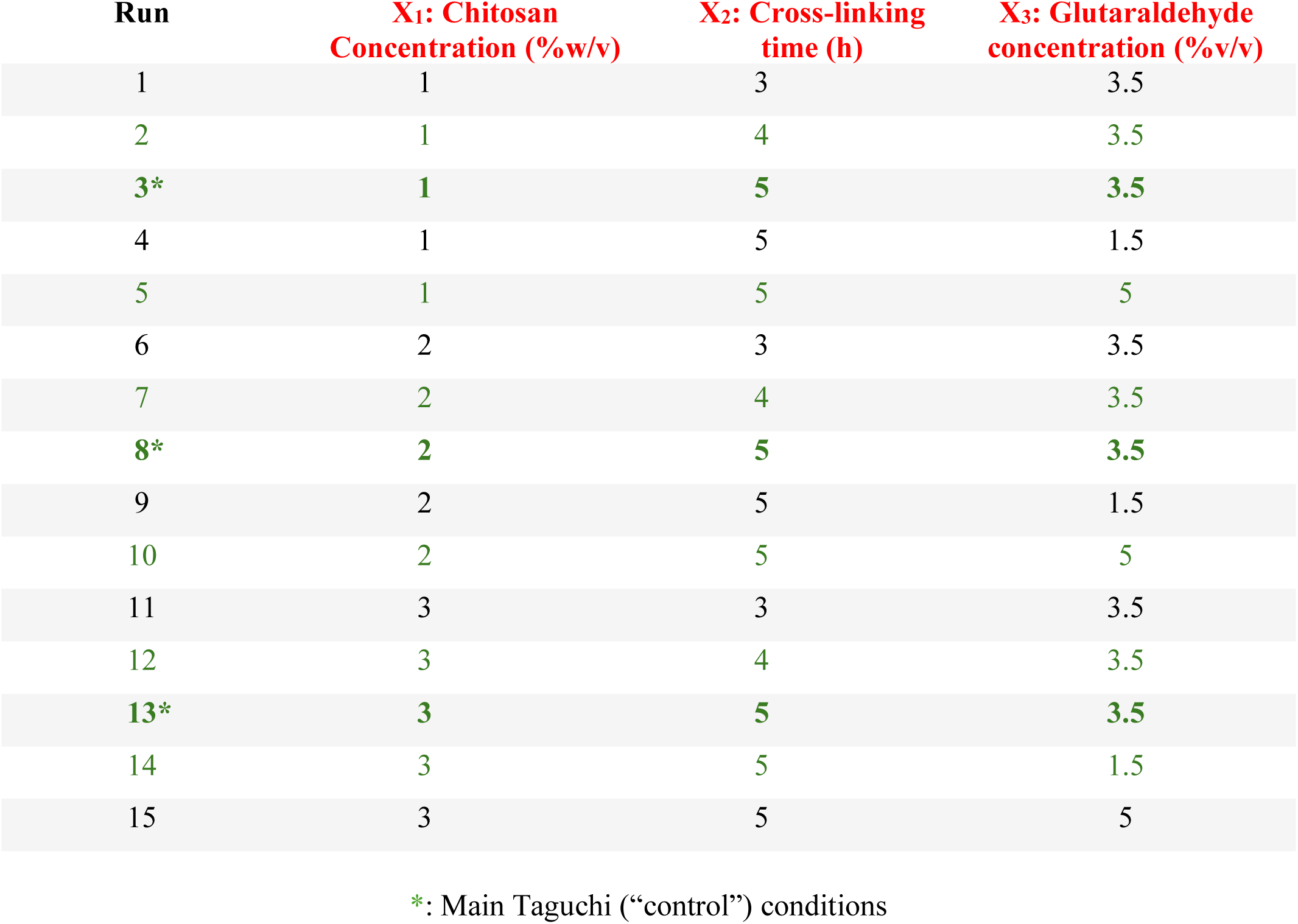
Changing conditions of drug loaded chitosan microspheres, obtained and designed from several preliminary experiments, for the optimization of size and encapsulation efficiency using Taguchi Orthogonal Design. Green marked conditions are taken as input factors levels for L_9_ orthogonal array of Taguchi design, while red marked X_1_, X_2_, and X_3_ are considered as factors.

| Run | X<sub>1</sub>: Chitosan<br>Concentration (%w/v) | X<sub>2</sub>: Cross-linking<br>time (h) | X<sub>3</sub>: Glutaraldehyde<br>concentration (%v/v) |
| --- | --- | --- | --- |
| 1 | 1 | 3 | 3.5 |
| 2 | 1 | 4 | 3.5 |
| 3* | 1 | 5 | 3.5 |
| 4 | 1 | 5 | 1.5 |
| 5 | 1 | 5 | 5 |
| 6 | 2 | 3 | 3.5 |
| 7 | 2 | 4 | 3.5 |
| 8* | 2 | 5 | 3.5 |
| 9 | 2 | 5 | 1.5 |
| 10 | 2 | 5 | 5 |
| 11 | 3 | 3 | 3.5 |
| 12 | 3 | 4 | 3.5 |
| 13* | 3 | 5 | 3.5 |
| 14 | 3 | 5 | 1.5 |
| 15 | 3 | 5 | 5 |

\*: Main Taguchi (“control”) conditions

### 2.7 Signal-to-Noise (S/N) Ratio Calculation

We calculated Taguchi S/N ratios on the nine core L₉ runs and other 6 extra conditions using the “nominal-the-best” metric for particle size of 6 μm and the “larger-the-better” metric for EE as we aim to have highest EE%. All S/N computations and main-effect plots were performed in Python (v3.10) using NumPy and Matplotlib. This allowed us to rank each condition both by mean performance and by robustness to noise. The full formulas are the following (n=2):

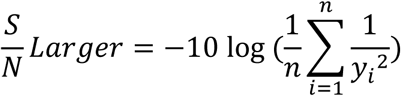

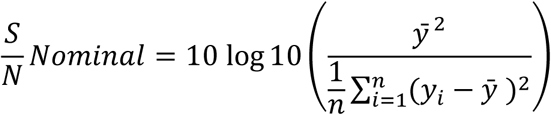

where;

- y_i_ is the i^th^ observed response,
- 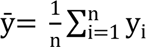 is the mean response for that condition,
- and n is the number of replicates under that condition.

### 2.8 Statistical Correlations and Machine-Learning-based Model Validation

To dig deeper into how each formulation parameter drives microsphere performance, both Pearson and Spearman correlation coefficients (Graphpad prism, v10.0.1) were computed on the full 18-averaged-point dataset (30 unique runs leading 15 averaged-point dataset + 3 replicates). Then, two types of predictive models on the full dataset were used: a tree-based Gradient Boosting Machine (using scikit-learn’s GradientBoostingRegressor with 100 trees and default learning rate) and a classical second-order polynomial regression (expanding [CS], [GA], and Time into all squared and pairwise interaction terms via ridge-regularized Poly² within a nested scheme). To ensure robust evaluation on this modestly sized dataset, each model was subjected to LOOCV to yield unbiased LOO-R² scores, and permutation-importance testing (1,000 shuffles per feature) to guard against overfitting and quantify each input’s marginal contribution.

Building on these validated models, we employed SHAP (v0.41.0) to translate otherwise “black-box” predictions into transparent, per-feature effect sizes. Using TreeExplainer for the GBM and LinearExplainer for the polynomial regression, we decomposed every prediction into additive contributions from [CS], [GA], Time, and their interactions.

### 2.9 Quadratic Response-Surface Modeling

Building on the validated polynomial model (Poly^2^) from section 2.8, we re-fit unregularized Poly^2^ fit on all 18 averaged-points to obtain explicit regression coefficients (β’s and γ’s) for main effects ([CS], [GA], Time), squared terms (CS², GA², Time²), and two-way interactions (CS·GA, CS·Time, GA·Time). To assess which of these terms contribute significantly to response variance, an analysis of variance (ANOVA) was performed on the fitted quadratic model in Python using the statsmodels library: the model was refit with “statsmodels.api.OLS,” and then “statsmodels.stats.anova.anova_lm” was used to decompose the total sum of squares into contributions from each main effect, squared term, and two-way interaction, yielding F-values and p-values for every coefficient. A threshold of p < 0.05 was used to define significance. We then generated a 60×60 meshgrid of [CS] (1.0–3.0 w/v%) and [GA] (1.5–5.0 v/v%) at fixed crosslinking time = 5h, transformed it via the same “PolynomialFeatures” instance, and predicted surface heights with “model.predict”. Filled contour and 3D surface plots were produced in Matplotlib.

It should be noted that we used two complementary Poly^2^ models built on the same polynomial basis (main effects, squares, and pairwise interactions). For predictive assessment, we used a ridge-regularized Poly^2^ within a nested scheme (outer LOOCV; inner 5-fold CV to select α via RidgeCV) and reported LOOCV R^2^; for inference and visualization (coefficients, p-values, contour/3D maps), we refit the same Poly^2^ with ordinary least squares (OLS) and performed ANOVA.

### 2.10 Statistical Analysis

We used one-way ANOVA followed by Tukey’s post-hoc test in GraphPad Prism software (version 10.0.1) to analyze our experimental data. A p-value of ≤ 0.05 was considered statistically significant. Significance levels are denoted as follows: n.s. (p > 0.05), * (p ≤ 0.05), ** (p ≤ 0.01), *** (p ≤ 0.001), and **** (p ≤ 0.0001).

## 3. RESULTS AND DISCUSSION

### 3.1 Characterization of DOX-Loaded Chitosan Microspheres

#### 3.1.1 FTIR

Fourier Transform Infrared spectra of chitosan, DOX, blank, and loaded microspheres were analyzed to identify characteristic functional groups and to demonstrate the insights into their molecular structures (**Figure 2a**). CS spectra revealed peaks corresponding to OH/NH stretching, which is the characteristic bands seen in 3358 and 3355 cm^−1^, CH stretching between 2873 and 2871 cm^−1^, and primary amine bending (–NH_2_) in around 1587 to 1560 cm^−1^. Doxorubicin displayed signature peaks at 3505 cm^−1^, 1625 cm^−1^, and 1507 cm^−1^ which are indicative of OH stretching and aromatic C=C stretching. The spectra of loaded microspheres revealed subtle shifts in peak positions at 3466, 2391, 2078, 1637 and 1281 cm^−1^, compared to the individual components. These shifts are attributed to intermolecular interactions, including hydrogen bonding and chemical interactions, between CS, GA, and DOX within the microsphere matrix. Hydrogen bonding between the hydroxyl and amino groups of CS and the carbonyl groups of DOX, as well as covalent interactions between the positively charged amino groups of CS and the negatively charged groups of GA, likely contribute to the observed spectral changes. These intermolecular interactions play a crucial role in stabilizing the microsphere structure and likely influencing the release kinetics of the encapsulated drug.

**Figure 2.**
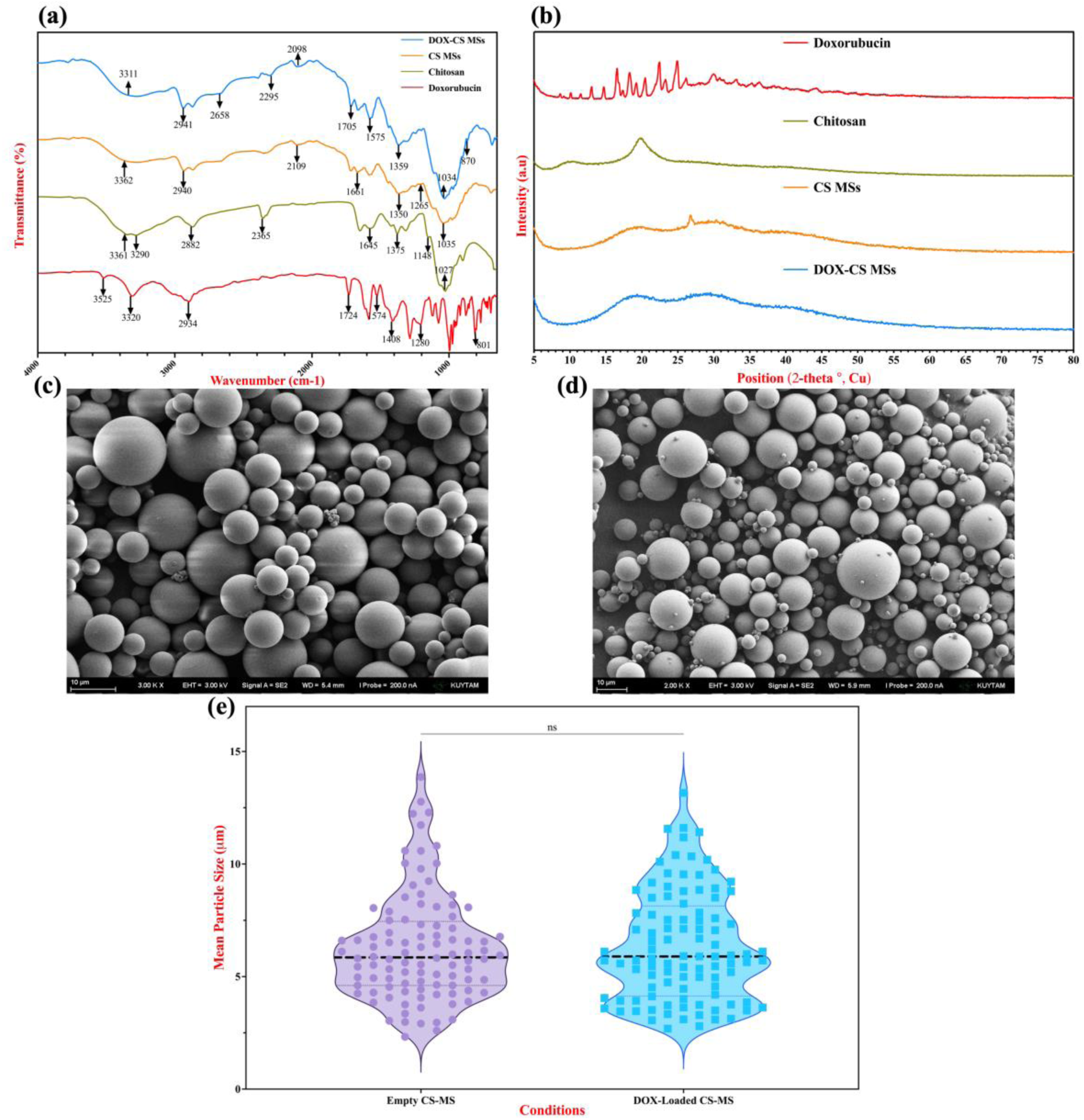
Physicochemical characterization and morphology of empty and DOX-loaded chitosan microspheres. **(a)** FTIR spectra of free doxorubicin (red), pristine chitosan polymer (olive), empty CS-MS (orange) and DOX-CS-MS (blue). **(b)** X-ray diffraction patterns of doxorubicin (red), chitosan (olive), empty CS-MS (orange) and DOX-CS-MS (blue). **(c, d)** Representative FE-SEM micrographs of (**c**) empty and (**d**) DOX-CS-MS showing smooth, spherical particles with uniform size. Scale bars = 10 μm. **(e)** Violin-box plot of particle-size distributions (n = 100) for empty (purple) and DOX-loaded (cyan) CS-MSs. The violin shape illustrates the kernel-density estimate; the central box spans the interquartile range, the black dashed line indicates the mean, and individual data points are overlaid. An unpaired two-tailed t-test showed no significant difference in mean diameter between groups (p > 0.05).

#### 3.1.2 XRD

**Figure 2b** presents the X-ray diffraction (XRD) patterns of Doxorubicin, CS, blank microspheres, and DOX-loaded CS MSs. Doxorubicin exhibits distinct crystalline peaks between 8° to 35° degree in 2θ range, confirming its highly crystalline nature. In contrast, pure chitosan and blank microspheres display broad diffraction patterns, indicative of an amorphous or semi-crystalline structure. On the other hand, the presence of low-intensity peaks corresponding to DOX in the loaded microsphere spectrum either suggests that DOX is in an amorphous state or successful DOX encapsulation within the polymer matrix. In other words, the reduction in crystallinity of chitosan after microparticle formation and further after DOX loading proves uniform distribution of DOX within the chitosan matrix. The altered peak intensities may be attributed to intermolecular interactions, including hydrogen bonding and van der Waals forces, between chitosan and doxorubicin during microsphere production.

#### 3.1.3 FE-SEM

FE-SEM analyses for each formulated CS MS and DOX-CS MSs were utilized to investigate nanoparticle size distribution, morphology, and surface characterization. **Figure 2c** and **2d** depicts the morphological examination of both optimized empty and DOX-loaded CS MSs, respectively, which display a narrowly distributed spherical shape with a smooth surface. According to observed FE-SEM images of various formulations, the homogeneity of the microspheres’ size seemed to be acceptable (data not shown). However, to obtain accurate findings related with the changing size among various formulations of DOX-loaded CS MSs and between the empty and DOX-loaded CS MSs, we have employed a detailed size investigation from different FE-SEM images and used Image J software to record at least 100 particles’ size. Recorded particle sizes were then analyzed further with frequency distribution, fitted into Gaussian Distribution model and analyzed with nonlinear regression to obtain final mean size data of each formulated microsphere. According to this analysis, the mean particle size of the empty and DOX-loaded CS MSs has been found as 5.693 and 5.822 μm, respectively, showing no significant difference between mean diameters. (**Figure 2e**). The resulting findings indicated that the incorporation of the drug did not significantly change the size of the microspheres.

### 3.2 Taguchi Screening Results

In this study, CS MSs were fabricated using a water-in-oil emulsion cross-linking approach in which the primary amines of CS are covalently bridged by GA. Briefly, an aqueous CS solution was emulsified into a mineral oil phase containing Span 80, and, after droplet stabilization at 1000 RPM mechanical stirring (selected based on preliminary SEM studies; **Figure S2**), GA was introduced to trigger cross-linking and microsphere solidification. Our preliminary optimization studies also revealed that using mechanical stirring at 1000 RPM consistently produced microspheres below 10 µm in diameter with our synthesis method, so this emulsification condition was held constant in all subsequent experiments (**Figure S3**).

Prior work in literature has shown that both emulsion droplet size and cross-link density govern final particle dimensions ^34^ while stirring speed and shear modality critically affect droplet breakup and drug retention ^35, 36^. However, the true interplay among crucial parameters such as biomaterial concentration, crosslinker level, and reaction time in determining micro or nanoparticles’ size and EE% remains still poorly understood, as these factors exhibit non-linear and synergistic effects that cannot be captured by one-variable-at-a-time (OVAT) approaches. Indeed, Wall *et al.* recently highlighted that, even in 2025, many chemical formulation studies continue to rely on OVAT screening rather than adopting efficient DoE methodologies, thereby incurring unnecessary experimental costs and delaying optimization processes ^14^.

Thus, to systematically map significant effects as a case-study, we designed 9 distinct formulations by varying three factors on three levels—CS concentration (1, 2, 3 % w/v), GA concentration (1.5, 3.5, 5 % v/v), and cross-linking time (3, 4, 5 h)—across a Taguchi L_9_ orthogonal array and also included six supplementary conditions chosen to fill gaps in the design space and enable reproducibility checks. This balanced yet expanded experimental set captures both the primary Taguchi combinations and additional edge- and center-point formulations, laying the groundwork for robust statistical analysis of resulting microsphere size and EE% of DOX incorporation.

To visualize our raw experimental data, a heat map of the normalized mean particle size, EE and DL for all 15 formulations were produced (**Figure 3a**). Each response variable was normalized within its row to a 0–1 range to highlight within-formulation trade-offs. Several formulations (notably F8, F10 and F13) demonstrated strong simultaneous performance across all three metrics. In contrast, certain formulations such as F7 achieved ideal EE but relatively modest size, highlighting the difficulty of optimizing multiple interdependent responses simultaneously. Statistical significance analysis via one-way ANOVA followed by Dunnett’s post hoc test (main conditions as control) indicated that many of the additional (non-Taguchi) formulations yielded comparable outcomes with main conditions selected. Interestingly, experimental results of DL% showed no consistent trends across the formulation space. Because DL is calculated relative to the total microsphere mass, which itself varies with particle size, yield and losses during washing, small changes in recovery or aggregation can disproportionately affect DL%, obscuring any direct parametric dependencies. Thus, we focus our predictive modeling and optimization primarily on size and EE, while reporting DL% here for completeness despite its sensitivity to mass-based variability.

**Figure 3.**
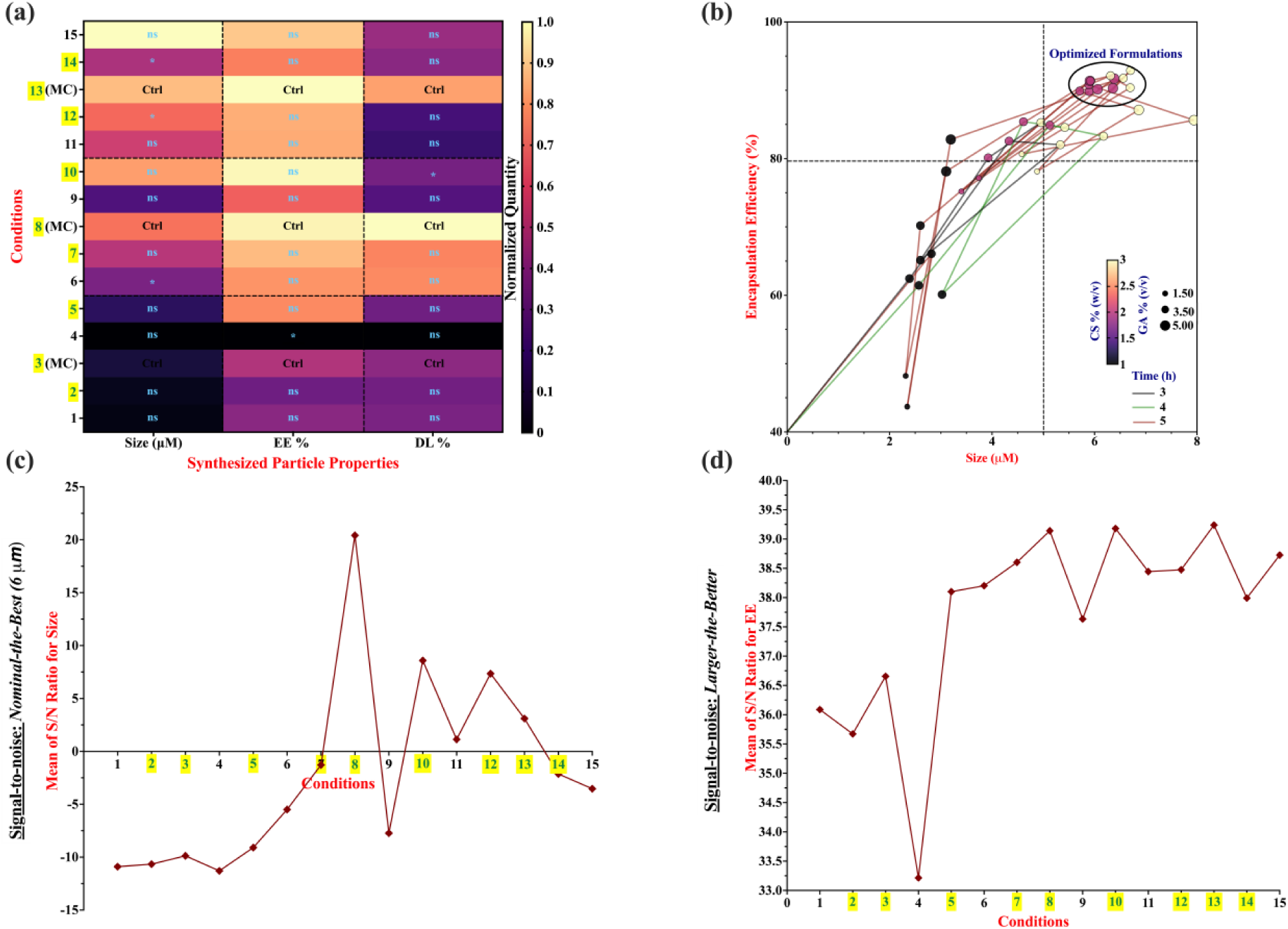
**(a)** Heat map of normalized mean responses for each condition (rows) and metric (columns). Responses—mean particle size (µm), encapsulation efficiency (EE, %) and drug loading (DL, %)—were scaled 0–1 within each row (0 = smallest/lowest observed mean; 1 = largest/highest observed mean). “*” indicates p < 0.05 versus the within-group control condition (the main Taguchi runs for each [CS]%) by one-way ANOVA with Dunnett’s multiple comparisons. **(b)** Bubble-cloud plot of all determined particle size versus EE% data for all experimental conditions including their replicates. Bubble color denotes [CS] (3 w/v%= yellow, 2%= red, 1%= black); connecting line color denotes crosslinking time (black = 3h, green = 4h, red = 5h); bubble size represents [GA] (smaller to bigger from 1.5 v/v% to 5.0%). Dashed black lines at size = 5 µm and EE = 80% mark target specifications, and the data-ellipse highlights the top-performing (“optimized”) formulations. **(c)** Signal-to-noise (S/N) ratios calculated using the “nominal-the-best” criterion for DOX-CS MS particle size. Each point represents one of the 15 experimental conditions. Taguchi-designed conditions (conditions 2, 3, 5, 7, 8, 10, 12, 13, 14) are highlighted in yellow. (**d)** Signal-to-noise (S/N) ratios for EE calculated using the “larger-the-better” criterion. Condition labels and color coding are identical to panel (**c**).

In **Figure 3b**, the relationship between particle size and EE is visualized through a multivariate bubble plot. Each data point represents a replicate of formulation, with size and color encoding synthesis time, CS concentration, and GA level, respectively. This visualization helps identify formulations that meet dual-target criteria — namely, a particle size in the range of 5–7 µm and EE exceeding 80% (denoted by dashed reference lines). The top-performing formulations (e.g. F8, F10, F13) clustered in the optimal upper-right quadrant, suggesting a favorable balance between structural compactness and drug EE. These findings are particularly important in light of literature showing that microparticles with tunable size and shape can be optimized for diverse biomedical applications, such as enhancing uptake by phagocytic cells for improved intracellular drug delivery, or enabling precise immune targeting ^1, 3^.

To quantitatively differentiate Taguchi-based formulations from supplementary runs and assess the exact effects of formulations, **Figure 3c** and **Figure 3d** present calculated S/N ratios using Taguchi robustness metrics. For particle size (**Figure 3c**), the “nominal-the-best” criterion was applied due to the defined optimal range, while EE (**Figure 3d**) followed the “larger-the-better” approach. S/N ratios were derived from replicate data and represent each formulation’s robustness against variability, a critical consideration in pharmaceutical applications where batch consistency is essential ^19, 25^.

In the S/N plot for size, only a subset of Taguchi-designed conditions demonstrated favorable values, suggesting that a smaller fraction of these designs successfully achieved tight particle size distributions. For EE, most Taguchi conditions displayed high and relatively uniform S/N ratios, affirming the utility of the Taguchi design in achieving robust drug encapsulation performance across synthesis variables. Importantly, these findings indicate that while Taguchi designs can be used for parameter screening, complementary out-of-array formulations may be essential to capture nonlinear interactions and optimize all performance metrics simultaneously, an observation consistent with recent reports advocating for hybrid DoE-ML frameworks in biomaterials optimization ^37^.

Collectively, the results presented in **Figure 3** demonstrate that the L_9_ Taguchi design was effective in narrowing the experimental parameter space and identifying several high-performing formulations with desirable particle sizes and encapsulation efficiencies. The integration of S/N ratio analysis added an important layer of robustness evaluation, enabling the assessment of response stability across replicates and highlighting formulations resilient to experimental noise.

However, as stated earlier, while the Taguchi method and its S/N-based evaluation offer a convenient metric for comparing overall performance, it remains limited in resolving the nuanced effects of individual parameters, particularly in systems governed by nonlinear interactions or higher-order dependencies. This constraint complicates mechanistic interpretation and restricts the predictive capacity of the optimization framework. To overcome these limitations and extract deeper insight into the influence of formulation variables on particle characteristics, we next employed supervised machine learning and statistical modeling approaches. These tools enable the construction of data-driven response surfaces, quantify variable importance, and facilitate predictive modeling across broader synthesis conditions, thereby enhancing both interpretability and translatability of the experimental findings.

### 3.3 Extended Design & Data-Driven Modeling

Building upon the initial screening performed, we validated experimental trends and uncover underlying relationships between formulation inputs and particle performance by applying both statistical correlation analysis and ML methods, ultimately enabling the identification of both linear and nonlinear dependencies across the formulation landscape. When training our statistical and ML models, we used one averaged data point per unique formulation rather than feeding in every individual replicate. Specifically, each of the 15 distinct [CS, GA, Time] combinations was performed in duplicate (plus a third repeat for our three “selected” Taguchi runs, making 18-data point), and the two measured Size and EE values per formulation were averaged before modeling. Our approach herein aimed to reduce redundant noise from replicate variability while preserving the core experimental trends, resulting in a cleaner training set for all analyses.

We first conducted a comprehensive correlation analysis using both Pearson’s (parametric) and Spearman’s (non-parametric) coefficients to interrogate linear and monotonic relationships between formulation parameters and microsphere characteristics (**Figure 4a**). This dual analysis revealed a strong positive correlation between CS concentration and particle size (Pearson *r* = 0.81; Spearman *rₛ* = 0.76), experimentally validating the size-regulating influence of polymer content. We think that, this trend is mechanistically attributed to the increased viscosity of the aqueous phase at higher CS concentrations, which impedes droplet breakup during emulsification, resulting in larger emulsion templates and, consequently, larger crosslinked particles. This viscosity-induced droplet retention also likely contributes to the enhanced encapsulation efficiency, reflected in a moderately strong correlation between CS concentration and EE% (*r* = 0.67; *rₛ* = 0.62). Denser matrices formed under higher CS content presumably minimize DOX loss during organic solvent washing (e.g., hexane, acetone), improving retention within the microspheres ^3, 34^.

**Figure 4.**
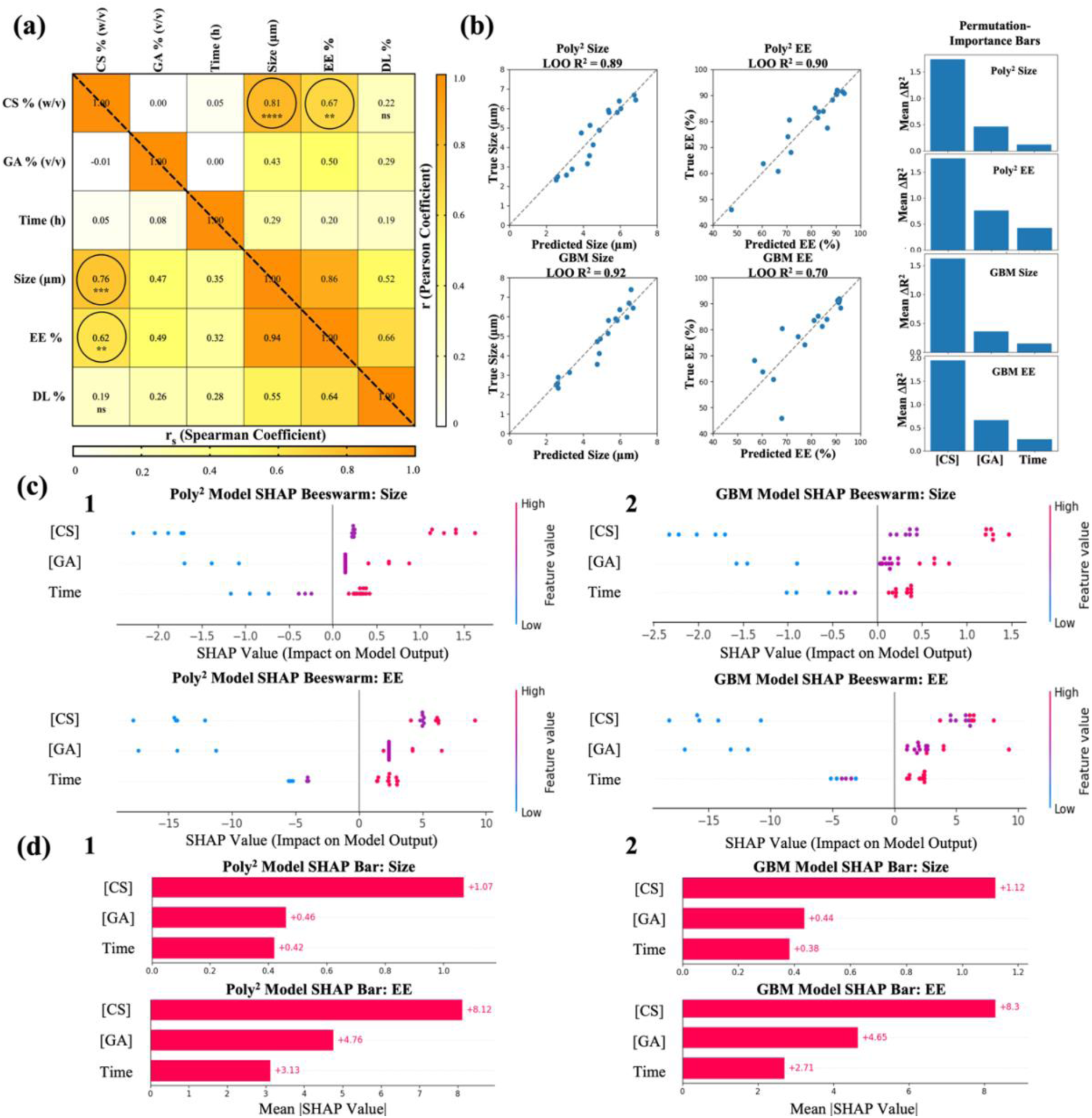
Comparative model evaluation and explainability for microsphere size and encapsulation efficiency. **(a)** Combined Pearson (upper triangle) and Spearman (lower triangle) correlation matrix between formulation factors—chitosan concentration (CS % w/v), glutaraldehyde concentration (GA % v/v), reaction time (h)—and measured particle properties (size, encapsulation efficiency [EE %], drug loading [DL %]). Significance levels: p < 0.05 (*), p < 0.01 (**), p < 0.001 (***), p < 0.001 (****), ns = not significant. **(b)** Leave-one-out cross-validation “predicted vs true” scatter plots for Size (µm) and EE (%), shown for the 2nd-order polynomial (Poly²) model (top row) and the Gradient Boosting Machine (GBM) model (middle row). Each subplot reports its LOOCV R². To the far right are permutation-importance bar charts (mean ΔR² when each feature is shuffled) for Poly² (upper two bars) and GBM (lower two bars), separately for Size (left) and EE (right). (c) SHAP beeswarm summary plots of per-sample SHAP values for each feature, for Size (upper panels) and EE (lower panels). Poly² results are given on (1), GBM on the (2). Dots are colored by the raw feature value (blue = low, red = high). **(d)** Corresponding mean |SHAP| bar charts (mean absolute SHAP values) for Size (top panels) and EE (bottom panels), again (1) for Poly² and (2) for GBM model. Numerical annotations at the end of each bar indicate the mean |SHAP|.

Similarly, the concentration of the crosslinker, GA, exhibited moderate correlations with both size (*r* = 0.43; *rₛ*= 0.47) and EE% (*r* = 0.50; *rₛ*= 0.49), suggesting its role in further reinforcing the matrix architecture. These correlations likely reflect the enhanced covalent bonding between aldehyde groups of GA and chitosan amines, which stabilizes particle morphology and encapsulated content during the post-synthesis purification steps. Notably, our experimental observations during the washing phase supported this interpretation, where formulations synthesized with higher GA content demonstrated visibly lower drug loss, consistent with tighter matrix formation ^23, 38^.

Reaction time showed a weaker but non-negligible influence on particle size and EE% with Spearman’s correlation (*rₛ* = 0.35 and *rₛ* = 0.32, respectively), suggesting that prolonged reaction kinetics may marginally support further matrix maturation or secondary crosslinking but are less dominant compared to CS or GA concentrations. As expected, drug loading percentage displayed poor correlation with all three input parameters, underlining its complex dependence on both particle architecture and drug–polymer interactions ^36^.

Interestingly, we also observed a strong interdependence between the two primary output variables, size and EE% (*r* = 0.86; *rₛ* = 0.94), indicating that formulations yielding larger particles also tended to encapsulate more DOX. This finding further reinforces the hypothesis that droplet integrity and internal matrix density—both strongly influenced by CS content—are pivotal in controlling DOX retention. In essence, particles synthesized under higher CS concentrations exhibit not only increased physical dimensions but also augmented internal capacity to entrap and preserve chemotherapeutic payloads ^9^.

Following the correlation analysis, we employed predictive modeling to quantitatively characterize the relationship between formulation parameters and resulting microsphere properties. To this end, we selected two complementary approaches tailored to the structure and size of our dataset. First, we used a second-order polynomial regression model (Poly²), which is widely adopted in the analysis of designed experiments for its simplicity and capacity to capture linear, quadratic, and two-factor interaction effects ^28^. Second-order models are particularly effective for low-to-moderate dimensional design spaces, where they can sufficiently approximate the true response surface without major overfitting ^39^. As a more flexible counterpart, we also implemented a Gradient Boosting Machine (GBM), a tree-based ensemble learning method well-suited for modeling complex, non-additive, and nonlinear relationships in small-to-moderate experimental datasets. By comparing these two models, we aimed to evaluate whether a basic polynomial regression could adequately describe our system or if the inherent interactions required a more powerful, data-driven approach. Together, these models enable both interpretability and predictive accuracy in understanding how formulation variables influence microsphere characteristics.

First of all, to quantitatively evaluate the predictive power of both models and minimize overfitting, particularly critical given the relatively small experimental dataset (n = 18), we applied LOO-CV. This approach systematically excludes one sample at a time during training and evaluates prediction error on that held-out point, enabling robust estimation of generalization performance without sacrificing data efficiency ^40^. As shown in **Figure 4b**, our ridge-regularized Poly^2^ model yielded high predictive accuracy for both microsphere size (R² = 0.89) and EE (R² = 0.90), supporting the adequacy of second-order regression in modeling key interactions within our design space. Interestingly, the GBM model demonstrated even stronger performance for size prediction (R² = 0.92), likely due to its ability to capture nonlinear relationships and threshold-like behaviors not modeled explicitly by quadratic terms. However, for EE%, the GBM underperformed slightly (R² = 0.70), suggesting that while EE trends are partially nonlinear, they may still be sufficiently described by additive or pairwise interactions captured by the Poly² model.

To better understand the contribution of individual formulation parameters to these predictions, we computed permutation-based feature importance (**Figure 4b**, right). This technique quantifies how much the predictive power of a model decreases (as measured by the drop in R², or ΔR²) when the values of a single input variable are randomly shuffled, thereby disrupting its relationship with the outcome ^41^. Consistently across all outputs and models, CS% was the most influential variable, especially for particle size, corroborating both our experimental findings and correlation analysis. GA concentration showed moderate influence on both EE% and size, while reaction time had relatively lower importance, consistent with its lower Pearson/Spearman correlation coefficients. Notably, the stronger size prediction performance of GBM may stem from its flexible modeling of CS–GA interaction or CS-induced viscosity thresholds that affect droplet formation dynamics, which are better captured by nonlinear tree ensembles than by polynomial equations.

Taken together, these results confirm that second-order regression offers both accuracy and interpretability for formulation systems governed by additive and pairwise interactions, while GBM provides enhanced flexibility to capture non-additive, complex influences, particularly for size-critical regimes.

To gain deeper insight into the internal decision logic of the trained models and to uncover the precise contribution of each input feature across all predictions, we employed SHAP, a unified framework for model interpretability grounded in cooperative game theory. While SHAP is most commonly used in large-scale datasets, recent studies have shown its effectiveness even for small- to moderate-sized experimental designs when carefully interpreted ^42, 43, 44^. Initially, we considered applying data augmentation strategies to expand the dataset, but given the limited variability and the risk of introducing artificial biases, we ultimately chose to proceed with the original 18-point experimental dataset to preserve interpretability and fidelity to the physical system. SHAP was applied to both Poly² and GBM models to enable transparent, model-specific attribution of formulation parameters, CS concentration, GA concentration, and reaction time, to predictions of particle size and EE%.

As illustrated in **Figure 4c**, beeswarm SHAP plots visualize the per-sample impact of each feature on predicted outcomes, colored by actual feature values (red = high, blue = low). Across both models, CS concentration emerged as the most consistently impactful parameter on microsphere size, with higher CS values positively contributing to larger size outputs. This aligns with experimental rationale, as increased polymer viscosity leads to larger emulsion droplets and subsequently larger particles. For EE, both models also attributed strong influence to CS concentration, followed by GA concentration, while reaction time had a minor but non-negligible effect. The mean absolute SHAP value bar plots (**Figure 4d**) further confirmed these trends, quantifying the average contribution magnitude of each variable. Notably, the GBM and Poly² models demonstrated close agreement in variable ranking and effect directionality, reinforcing the robustness of the underlying relationships despite the models’ structural differences. Collectively, these SHAP results further validate that CS concentration is the dominant factor governing both physical and functional characteristics of the synthesized microspheres, while GA and time exhibit more subtle but context-specific influences.

In summary, the predictive modeling results from both the polynomial and GBM frameworks not only aligned well with our experimental findings (**Figure 3**), but also provided quantitative insight into the parametric influence on microsphere properties. The high fidelity between predicted and observed outcomes demonstrates that data-driven modeling can robustly validate and extend experimental screening results.

### 3.4 Predictive Model Construction and Surface Modelling

Observing the validated predictive capacity of two different models, Poly² was selected (LOO R² = 0.89) and EE (LOO R² = 0.90) to derive interpretable equations that quantitatively relate synthesis parameters to microsphere characteristics. In contrast to black-box models, the polynomial regression approach enables the explicit formulation of a mathematical relationship between input variable and output responses. This formulation is particularly advantageous for experimental systems, as it allows real-time predictions, intuitive optimization, and scalable implementation across different formulation ranges ^14^. Therefore, we re-fit identical Poly^2^ model with OLS to experimental data summarized in **Table 2**, then generated two- and three-dimensional response surfaces at the fixed crosslinking time of 5h, reflecting the most commonly used synthesis duration in our protocol. This dual-track Poly^2^ strategy preserves predictive rigor for performance reporting, while the OLS refit yields explicit equations and interpretable contour/3D maps that expose curvature and critical interaction zones, directly guiding factor selection to hit target particle size and EE.

**Table 2.** Experimental runs and responses for all formulations. Green runs show Taguchi conditions.

| Run | X<sub>1</sub>: CS% | X<sub>2</sub>: Time(h) | X<sub>3</sub>: GA% | Response 1:<br>Size<br>μm | Response 2:<br>EE<br>% |
| --- | --- | --- | --- | --- | --- |
| 1 | 1 | 3 | 3.5 | 2.495 | 63.785 |
| 2 | 1 | 4 | 3.5 | 2.5867 | 60.775 |
| 3* | 1 | 5 | 3.5 | 2.8835 | 68.14 |
| 4 | 1 | 5 | 1.5 | 2.329 | 45.95 |
| 5 | 1 | 5 | 5 | 3.15 | 80.465 |
| 6 | 2 | 3 | 3.5 | 4.1285 | 81.34 |
| 7 | 2 | 4 | 3.5 | 4.87 | 85.15 |
| 8* | 2 | 5 | 3.5 | 5.9055 | 90.585 |
| 9 | 2 | 5 | 1.5 | 3.5705 | 74.18 |
| 10 | 2 | 5 | 5 | 6.372 | 91.32 |
| 11 | 3 | 3 | 3.5 | 5.143 | 83.645 |
| 12 | 3 | 4 | 3.5 | 5.7975 | 83.915 |
| **13\*** | **3** | **5** | **3.5** | **6.6995** | **91.635** |
| 14 | 3 | 5 | 1.5 | 4.73 | 77.39 |
| 15 | 3 | 5 | 5 | 7.4025 | 88.372 |
| 16 † | 2 | 5 | 3.5 | 5.8 | 90.35 |
| 17 † | 3 | 5 | 3.5 | 6.437 | 91.947 |
| 18 † | 2 | 5 | 3.5 | 5.985 | 90.765 |

\* Main Taguchi (“control”) conditions
† Repeats of runs 8, 13, 10 for reproducibility checks, respectively.

After fitting full second-order polynomial equations to both particle size and EE, the measured responses are expressed as the following:

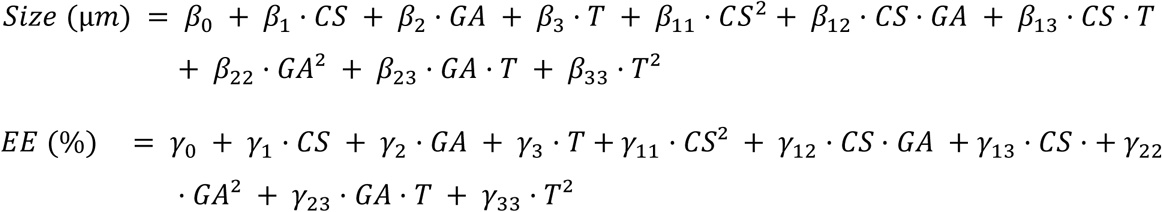

OLS fitting of Poly^2^ to our data produced the high coefficients of determination shown in **Table 3**, and yielded the individual regression coefficients for each main effect, squared term, and two-way interaction. We then subjected each full quadratic model to ANOVA to test which terms explain significant portions of response variance (**Table 4**). For particle size, both the linear CS term (F = 9.96, p = 0.012) and its squared term (F = 23.51, p = 0.001) are highly significant, as is the CS·GA interaction (F = 10.34, p = 0.011), confirming that polymer concentration, as well as its interplay with crosslinker, dominates size control. By contrast, GA and time alone (and their interactions with time) do not reach significance for size. Unlike the strong bivariate correlation we observed between GA% and particle size (r≈0.47 in **Figure 4a**), the linear GA term in our full quadratic model proved non-significant (F=0.29, p=0.60; **Table 4**). That apparent paradox simply reflects the fact that, once we “hold constant” CS concentration, cross-linking time, and all squared and interaction terms, the pure GA→Size slope carries little independent weight. In contrast, both the GA^2^ term (F=6.01, p=0.037) and the CS·GA interaction (F=10.34, p=0.011) emerge as significant contributors, confirming that GA primarily modulates size via non-linear and synergistic effects with [CS]. In other words, GA does matter, but its influence is captured not by a simple straight-line effect, but by the curvature and cross-term in the quadratic surface. This conclusion aligns perfectly with our SHAP analyses (**Figure 4c–d**), which ranked GA behind CS but still flagged it as a meaningful driver of both size and EE once interactions are taken into account.

**Table 3.** Fitted quadratic-regression coefficients for particle size (µm) and encapsulation efficiency (EE, %) as functions of CS concentration, GA concentration and crosslinking time. R² is the coefficient of determination for each full second-order model; “Intercept” and subsequent columns list the estimated regression coefficients (β’s and γ’s) for the main effects (CS, GA, Time), squared terms (CS², GA², Time²), and two-way interactions (CS·GA, CS·Time, GA·Time). Positive values indicate an increasing effect on the response; negative values indicate a decreasing effect.

| *Response* | *R<sup>2</sup>* | *Intercept* | *CS* | *GA* | *Time* | *CS<sup>2</sup>* |
| --- | --- | --- | --- | --- | --- | --- |
| *Size* | <u>0.983</u> | −0.569 | +2.571 | +0.670 | −0.940 | −0.692 |
| *EE* | <u>0.986</u> | +19.338 | +60.879 | +19.787 | −27.902 | −9.661 |
| *Response* | *CS·GA* | *CS·Time* | *GA<sup>2</sup>* | *GA·Time* | *Time<sup>2</sup>* |  |
| *Size* | +0.267 | +0.204 | −0.141 | +0.060 | +0.117 |  |
| *EE* | −3.258 | −0.107 | −2.133 | +1.277 | +3.429 |  |

**Table 4.** ANOVA summary for full quadratic model, sums of squares, F-values, and p-values for each main effect, squared term, and interaction in the Size and EE regressions.

| *Term* | *Size_Sum*<br>*Sq* | *F-value* | *p-value* | *EE_Sum*<br>*Sq* | *F-value* | *p-value* |
| --- | --- | --- | --- | --- | --- | --- |
| **CS** | 0.848 | 9.96 | 0.012 | 475.229 | 105.33 | <0.001 |
| **GA** | 0.025 | 0.29 | 0.603 | 54.556 | 12.09 | 0.007 |
| **Time** | 0.085 | 0.99 | 0.345 | 46.647 | 10.34 | 0.011 |
| **CS<sup>2</sup>** | 2.001 | 23.51 | 0.001 | 389.897 | 86.42 | <0.001 |
| **GA<sup>2</sup>** | 0.511 | 6.01 | 0.037 | 117.476 | 26.04 | 0.001 |
| **Time<sup>2</sup>** | 0.029 | 0.34 | 0.573 | 25.098 | 5.56 | 0.043 |
| **CS·GA** | 0.880 | 10.34 | 0.011 | 131.175 | 29.07 | <0.001 |
| **CS·Time** | 0.277 | 3.25 | 0.105 | 0.076 | 0.02 | 0.900 |
| **GA·Time** | 0.020 | 0.23 | 0.642 | 0.215 | 0.05 | 0.832 |
| **Residual** | **0.766** | — | — | **40.607** | — | — |

For EE, virtually every CS term (linear and quadratic, F = 105.33 and 86.42, both p < 0.001), GA (F = 12.09, p = 0.007), GA² (F = 26.04, p = 0.001), time (F = 10.34, p = 0.011) and the CS·GA interaction (F = 29.07, p < 0.001) significantly contribute to EE variance, whereas CS·Time and GA·Time remain non-significant. These ANOVA results again quantitatively validate our earlier SHAP and permutation-importance findings (**Figure 4**): CS concentration is the primary driver of both size and EE, GA content fine-tunes encapsulation, and crosslink density (via CS·GA) modulates both outputs, while time plays a secondary or insufficient role.

Finally, to translate our quadratic models into intuitive design maps, we held cross-linking time constant at 5h and plotted both contour and 3D response surfaces for particle size and EE (**Figure 5**). In panels (**a**) and (**b**), contour heatmaps show “iso-size” and “iso-EE” curves across CS and GA concentrations, with the red “×” marking our Taguchi runs along the heatmaps. Panels (**c**) and (**d**) present the same data as three-dimensional surfaces, confirming that larger CS and GA levels synergistically increase both size and encapsulation, but with diminishing returns at the highest extremes, exactly as suggested by our significant squared and interaction terms. These visualizations provide an actionable guide for selecting CS/GA combinations to target specific size or EE goals in future formulations of this microsphere system.

**Figure 5.**
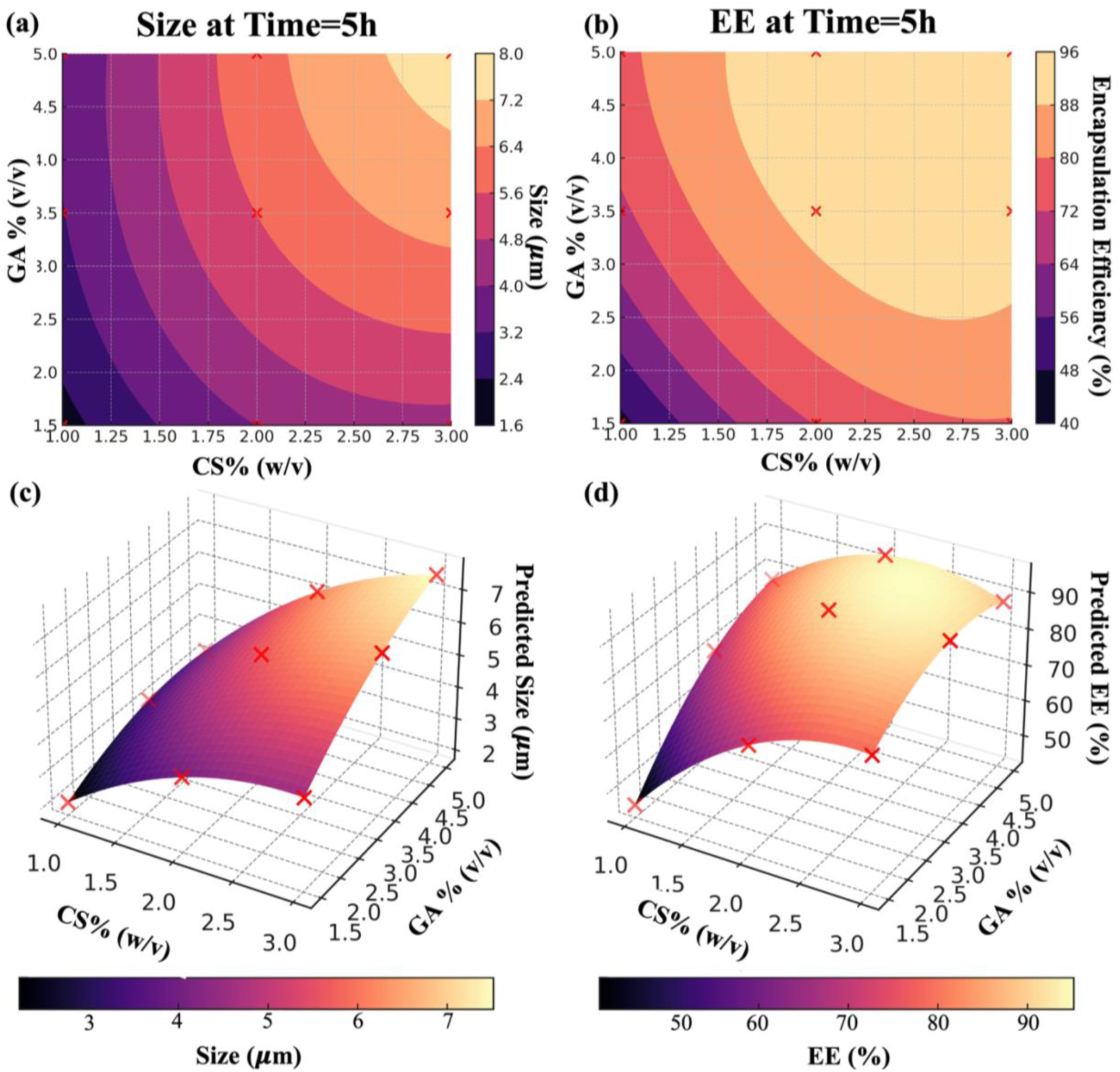
**(a)** Contour plot of predicted microsphere diameter (µm) versus CS and GA concentrations at fixed cross-linking time (5h). **(b)** Contour plot of predicted EE% under the same conditions. **(c,d)** Corresponding 3D response-surface plots for size and EE, respectively. Red “×” markers denote the nine main Taguchi experimental conditions.

Although we validated our findings through multiple complementary analyses, the core quadratic surface method seems to remain a highly accessible and reliable tool for capturing nonlinear effects and interactions in microsphere synthesis. Importantly, our workflow began with a simple Taguchi L_9_ screen to decrease the full parameter space, dramatically reducing experimental effort, and then seamlessly connected to richer data-driven modeling to uncover and quantify higher-order behavior. This hybrid strategy demonstrates that even in complex chemical systems, a classic response-surface framework can serve as both a practical and powerful guide for formulation optimization.

## 4. CONCLUSIONS

In this study, we present a systematic and data-driven framework to optimize a particulate drug-carrier system, using DOX-CS MSs as a proof of concept. We combine a Taguchi orthogonal design with statistical modeling and ML-based interpretability to study how CS concentration, GA content, and crosslinking time affect two key outputs: particle size and EE%. To our knowledge, this is the first application of a Taguchi–ML hybrid framework directly to a real experimental dataset for microsphere synthesis, delivering both mechanistic insight and predictive models.

We began with an L₉ Taguchi screen to narrow the design space and identify promising conditions. We then added targeted extra formulations to resolve nonlinear effects and interactions. Correlation analysis, polynomial regression, gradient-boosting models, and SHAP agreed on the main result: CS concentration is the dominant factor for both size and EE. GA shows secondary but synergistic effects, especially in combination with CS. Quadratic response-surface modeling (Poly²) with ANOVA confirmed these trends and produced explicit predictive equations. The models visualize size and EE landscapes for formulation targeting and achieved high in-sample accuracy (R² = 0.983 for size; R² = 0.986 for EE). Our hybrid workflow reduces experimental burden while preserving the ability to map factor influences quantitatively, making it applicable to other particulate systems with complex formulation–property relationships.

The methodology applied here offers several advantages: (**i**) the combination of Taguchi screening and ML-driven modeling enables rapid factor prioritization and detection of nonlinear interactions that traditional Taguchi interpretation alone may miss; (**ii**) the predictive surface models can be directly used for formulation targeting in similar chitosan systems or extended to other biopolymer-based micro-nanospheres; and (**iii**) the SHAP framework adds transparency to ML predictions, aiding mechanistic understanding. However, some limitations should be acknowledged. The relatively small dataset, although representative for proof-of-concept, constrains model generalizability beyond the tested factor ranges. Additionally, drug loading percentage (DL%) proved inconsistent across replicates, likely due to size-dependent yield variability, highlighting the need for improved mass recovery consistency in future studies.

Future work should extend this hybrid optimization framework to larger and more diverse experimental datasets with easier chemical synthesis methods, incorporate additional response variables such as release kinetics and cytocompatibility, and evaluate its utility in dual-drug or multi-stimuli-responsive microsphere systems. Collectively, our demonstrated approach provides a reproducible and generalizable blueprint for accelerating the optimization of complex particulate synthesis processes, bridging efficient experimental design with data-driven predictive modeling to streamline the path toward application-ready drug delivery carriers required for targeted biomedical needs.

## Data Availability Statement

The data/codes supporting this study’s findings are available from the corresponding author upon reasonable request.

## CRediT Authorship Contribution Statement

**D.D:** Writing – original draft, Writing – review & editing, Methodology, Investigation, Data curation, Visualization, Validation, Conceptualization. **S.K*:** Writing – review & editing, Writing – original draft, Supervision, Resources, Project administration, Funding acquisition, Conceptualization, Validation.

## Acknowledgements

The authors gratefully acknowledge the facilities and technical support provided by the Koç University Research Center for Surface Science (KUYTAM) and the Koç University Research Center for Translational Medicine (KUTTAM). D.D. also extends sincere appreciation to TÜBİTAK for the BIDEB scholarship support.

## Supporting Information

The Supporting Information is available free of charge.

