## Supplementary material for "Taguchi–Machine Learning Hybrid Framework for Optimization of Particulate Drug Delivery Systems": Figure S1, S2, S3

Doğukan Duymaz, Seda Kizilel*

Chemical and Biological Engineering, Koç University, Sarıyer, İstanbul, Türkiye

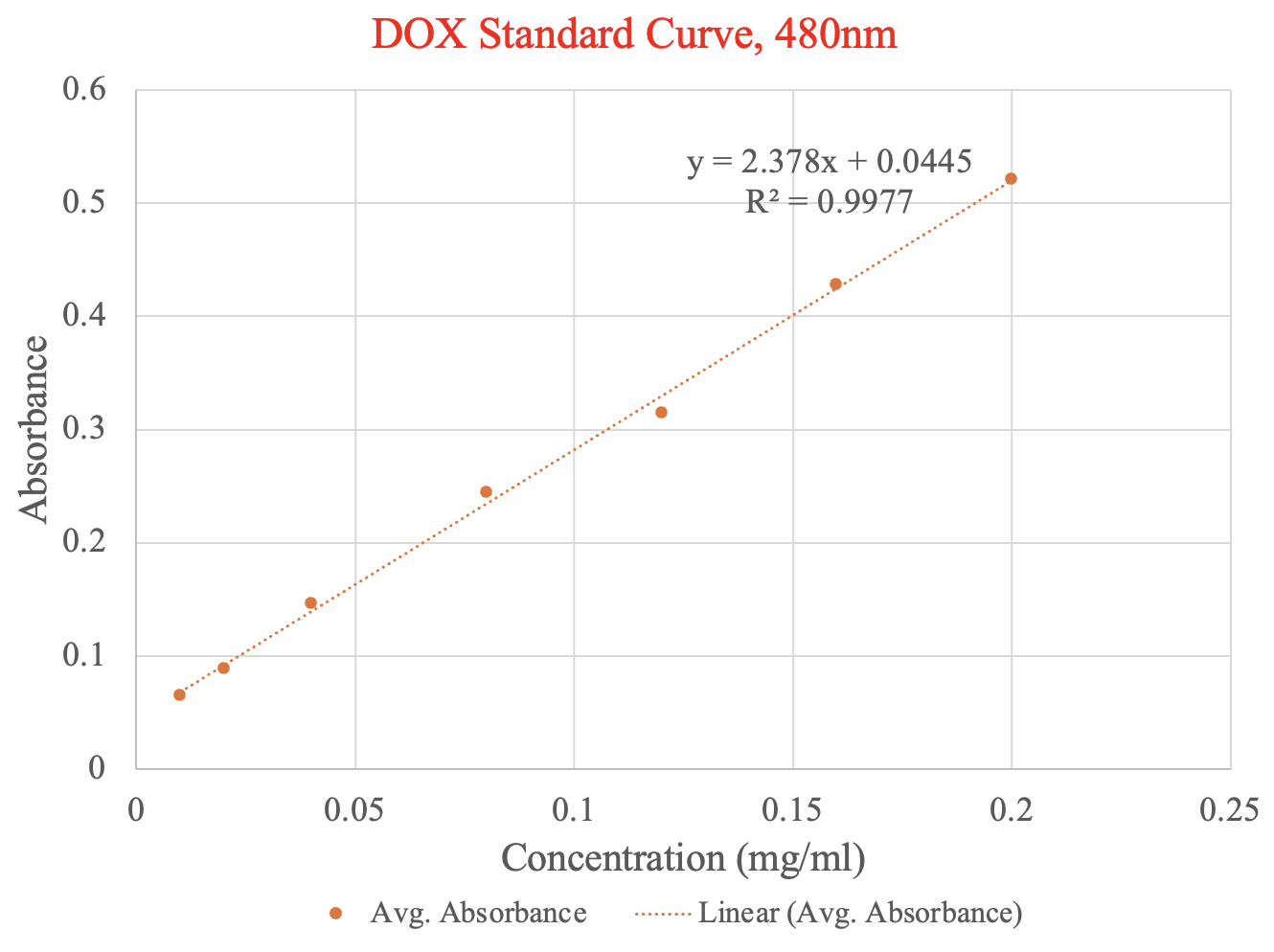

**Figure S1**. DOX standard calibration curve at 480 nm. A strong linear correlation (R^2^ = 0.9977) was obtained between absorbance and increasing DOX concentration (0–0.2 mg/ml).

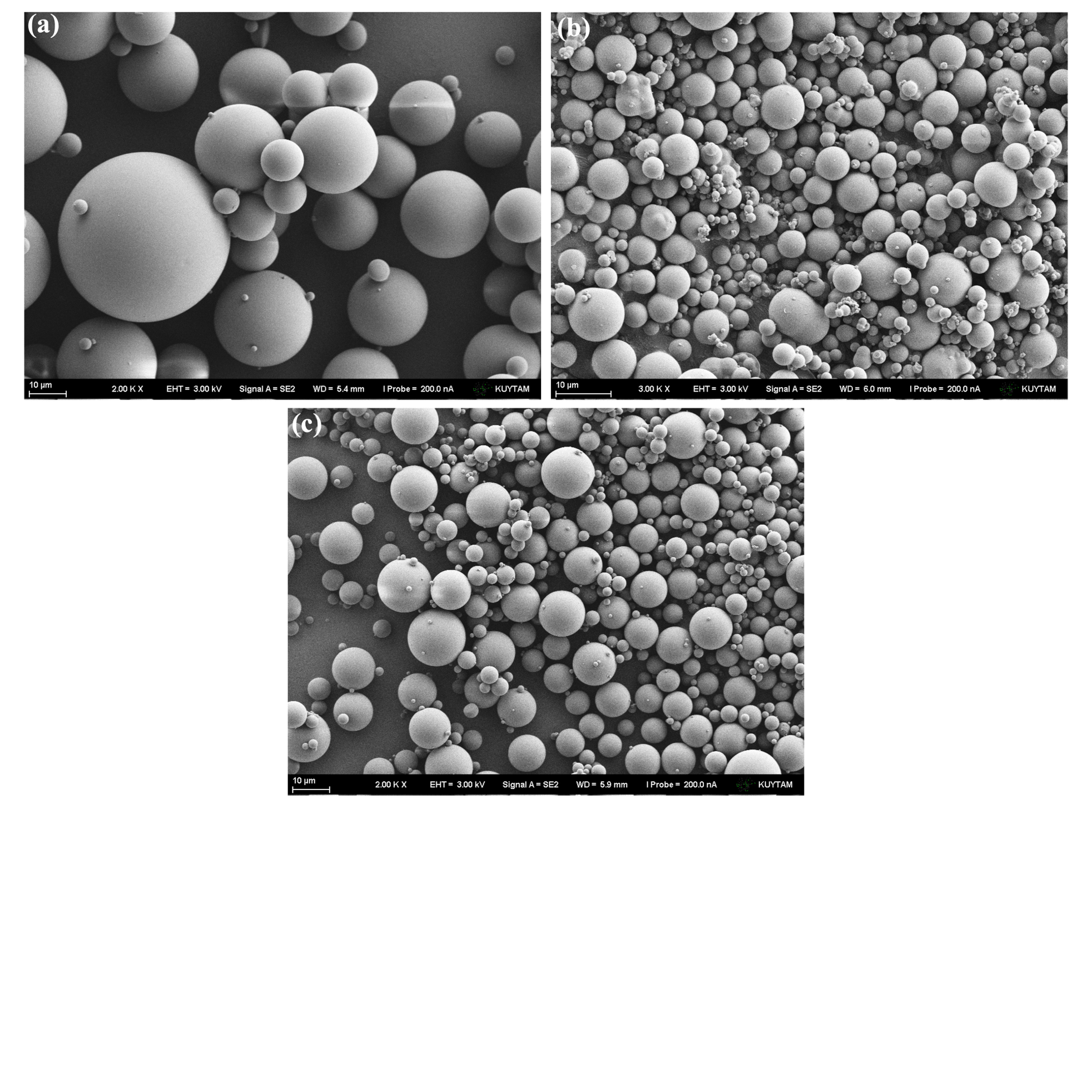
**Figure S2.** SEM images illustrating the effect of mechanical stirring speed on the morphology and size distribution of chitosan microspheres synthesized using the same protocol. (**a**) Stirring speed below 1000 RPM resulted in generally larger particles (> 10 µm) with smoother surfaces. (**b**) Stirring speed above 1000 RPM yielded smaller particles with increased heterogeneity. (**c**) Optimized stirring speed of 1000 RPM produced uniform, well-defined spherical particles. All scale bars: 10 µm.

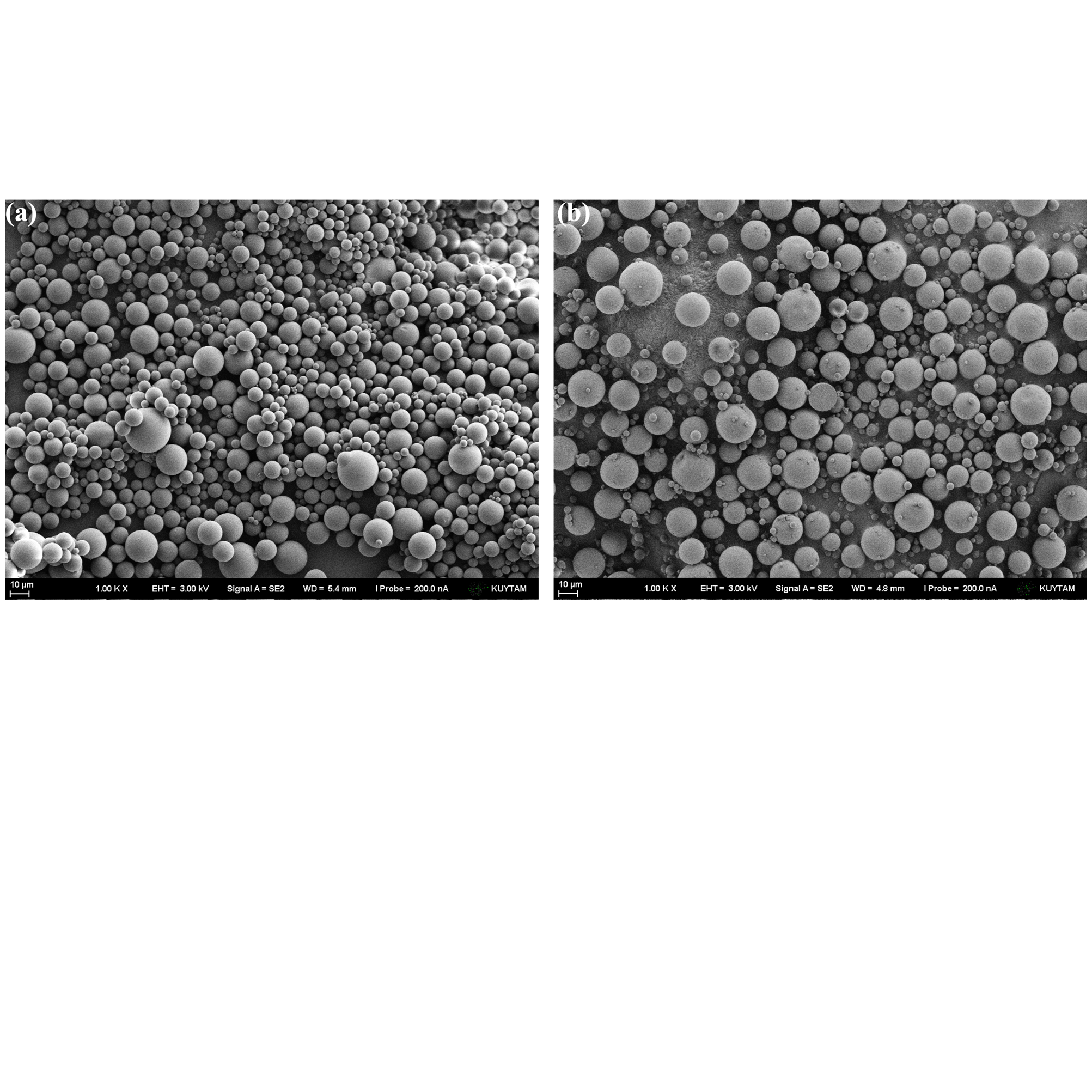
**Figure S3.** SEM images comparing the effect of stirring method on chitosan microsphere morphology using the same synthesis protocol. (**a**) Mechanical stirring at the optimized speed of 1000 RPM produced uniform, spherical particles with smooth surfaces. (**b**) Magnetic stirring under identical conditions resulted in less uniform morphology and occasional surface irregularities. All scale bars: 10 µm.
